# Artificial soil systems: A tool for investigating microbial life strategy effects on substrate mineralization

**DOI:** 10.1101/2025.04.04.647204

**Authors:** Kanade Fujiwara, Tomoyuki Makino, Toru Hamamoto

## Abstract

Soil microbes play a critical role in carbon (C) cycling, however, the influence of microbial life strategies and their interactions on C mineralization remains poorly understood. This study aimed to investigate how r- and K-strategist bacteria influence glucose mineralization using an artificial soil system, focusing on *Bacillus subtilis* and *Streptomyces cinnamoneus*. Through a 14-day incubation experiment, we found that *B. subtilis* exhibited rapid and high respiration rates, while *S. cinnamoneus* showed slower, delayed respiration rates, supporting their respective r/K classification. In co-culture treatments, cumulative glucose mineralization converged with *B. subtilis* monoculture levels and positively correlated with its relative abundance. These findings demonstrate that artificial soil systems effectively reveal how microbial interactions drive C dynamics, offering a controlled approach to elucidate mechanisms underlying soil C cycling.

## Introduction

One gram of soil can contain as many as 10 billion microbes (Torsvik & Øvreås, 2002), which play a crucial role in the soil carbon (C) cycle. Soil microbes decompose soil organic matter (SOM) to generate energy (i.e., ATP) for their metabolic activities, and assimilate it into their biomass while releasing CO_2_ through respiration. Microbial products (i.e., necromass, protein, and DNA) accounted for 10-80% of SOM (Chao et al., 2019), while microbial respiration accounts for more than 60% of soil CO_2_ emissions (Iven et al., 2023; Shang et al., 2024).

The r/K selection theory, proposed by Pianka (1970), has been widely applied to understand the relationship between soil microbial community and soil C dynamics (Chen et al., 2022; Cheng et al., 2022; Zheng et al., 2024). The relative abundance of r- and K-strategists significantly influences soil C mineralization rates. K-strategists are characterized by slow growth rates and are associated with enhanced soil C sequestration (Blagodatsky et al., 2010; Hu et al., 2023), while r-strategists, characterized by rapid growth rates, dominated soil with high SOC content and thus enhanced soil respiration (Blagodatsky et al., 2010; Shang et al., 2024). Recent studies reported that the microbial community shifted from K-strategists to r-strategists with increasing SOM and higher soil respiration in forest and agricultural soils (Li et al., 2023; Wang et al., 2021; Zheng et al., 2024).

Traditionally, soil microbes are classified into r- and K-strategists based on the correlation between C mineralization rates and microbial traits (e.g., Gram positive/negative and GC contents) or the relative abundance at the phylum level. For example, high GC content bacteria, such as Actinobacteria, are classified as K-strategists, while low GC content bacteria, such as Firmicutes, are categorized as r-strategists (Aliperti et al., 2023). In the classification based on Gram staining, contrastingly, gram-positive bacteria which includes Firmicutes and Actinobacteria are categorized as K-strategists (Sun et al.,2021). Previous meta-analytical studies showed that C mineralization rate positively correlated with Bacteroidetes abundance, suggesting r-strategists, while negatively correlated with Acidobacteria abundance, indicating K-strategists, while the phyla Firmicutes and Actinobacteria were not significantly correlated to the C mineralization rates (Fierer et al, 2007; Zhang et al., 2018; Yang et al., 2021). This study identifies the following key issues: (1) The mineralization rates of microbial species have not been directly evaluated in soil. (2) Consequently, the classification of r/K-strategy based on community assessments or traits in previous studies may lead to contradictory results depending on the experimental conditions. However, the underlying mechanism of interactions among r- and K-strategists, as well as the causal relationship between life-history strategies and C dynamics, remains unclear. A deeper understanding of C cycling through microbial regulation requires integrating data from individual microbes and simplified microbial communities with comprehensive analyses of terrestrial microbial ecosystems (Trivedi et al., 2013; Nazaries et al., 2013).

Liquid and agarose media are commonly used in microbial research to investigate interspecific interaction (Straight et al., 2006; Yan et al., 2021). However, Del Valle (2022) reported significant differences in C dynamics between liquid culture and natural soil conditions, suggesting that simplified systems might not accurately reflect microbial dynamics in natural soil environments. This highlights the need for experimental systems which can bridge the gap between simplified media and complex natural soils. Sterilized soil, including autoclaving and □-irradiation, offer one alternative approach to control microbial composition (Wertz et al., 2007; Delmont et al., 2014), while sterilization procedures can alter soil chemical or physical properties (Wolf et al., 1989; Ellis, 2004). In addition, some phyla (e.g., Firmicutes and Proteobacteria) are resistant to autoclaving (Li et al., 2019).

Artificial soil systems represent another alternative approach which has been employed to investigate the impact of abiotic factors on soil microbial C dynamics (Maeda et al., 2020; Rakhsh et al., 2020; Luiz A et al, 2020; Xu et al., 2024). Previous studies cultured single species (Guenet et al., 2011; Lennon et al., 2012) or simple microbial communities (Ellis, 2004; Wolf et al., 2013) in artificial soils. For instance, Ellis (2004) demonstrated that co-culturing *B*.*thuringinesis* with *P*.*fluorescens* in artificial soils reduced the mortality of B. thuringiensis by fourteen-fold compared to monoculture. Similarly, Wolf et al. (2013) showed that *B. weihenstephanensis* and *S*.*atratus* in the microcosms varying matric potential and pore size distribution. While these studies assessed microbial mortality in artificial soils, they did not evaluate how interspecific interactions affected substrate mineralization rates, leaving a significant gap in our understanding of how microbial interactions influence C dynamics.

The objective of this study was to investigate the role of different microbial species and their interactions in glucose mineralization using artificial soil system. We hypothesize that substrate mineralization differs based on microbial life strategies (r/K) and significantly influenced by interspecific interactions. To test these hypotheses, we conducted a 14-day incubation experiment with two microbial species in an artificial soil environment.

## Material & Method

### Artificial soil

Artificial soil components were developed based on previous reports by Ellis (2004) with minor modifications (**Table 1**). Briefly, quartz sand was used as the sand fraction, while kaolinite and Japanese acid clay were used for the clay fraction. In addition, calcium carbonate and humic acid were used as the buffering capacity and microbial nutrients, respectively. The soil pH was adjusted to 7.2. The total C and N contents of the artificial soils were 11.5 mg g□^1^ soil and 0.5 mg g□^1^ soil, respectively. The artificial soils were sterilized with an autoclave (121°C, 20min) and stored until use.

**Table 1.**
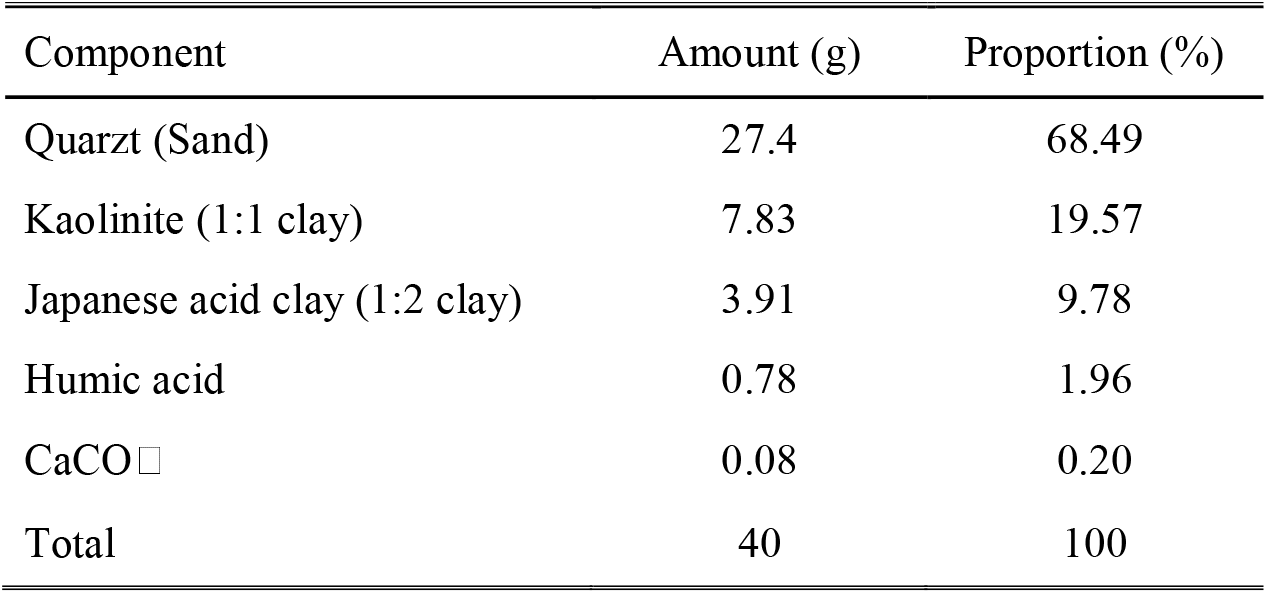
Composition of the artificial soil.

### Experimental design

Two bacterial strains were inoculated in the artificial soils: *Bacillus subtilis* (*B*.*subtillis*; NBRC101584), and *Streptomyces cinnamoneus* (*S. cinnamoneus*; NBRC13823). *Bacillus* is the genus of the phylum Firmicutes, and *Streptomyces* is the genus of the phylum Actinobacteria. Both genera are among the most abundant in natural soils and play key roles in soil organic matter decomposition (Nonthakaew et al., 2022; Saxena et al., 2020; Wolf et al., 2013). Both strains were recovered from L-dried ampoules according to the method described by manufacturer’s instructions and then stored at −80□ in glycerol stocks until the use. Prior to inoculation into the artificial soils, each bacteria was cultivated and maintained in growth phase at 30°C according to manufacturer’s instructions. The bacterial cultures were then centrifuged at 2150 × g for 10 minutes and washed with phosphate-bu□ered saline (PBS, pH 7.0) to remove medium components which could induce bacterial respiration (Baker et al., 2020; Jiang et al., 2023). Both bacterial strains were resuspended in PBS and adjusted to OD_600_ of 0.1. Additionally, DNA extraction was performed using the Mighty Prep reagent for DNA (Takara Bio, Inc., Japan), according to the manufacturer’s protocol. The concentration of extracted DNA was quantified by using the QuantiFluor dsDNA system (Promega, USA).

The microcosm consisting of 40 g (equivalent dry mass) of sterilized artificial soil component mixture was placed in a sterilized 50 mL bottle (**Fig. S1**). The experimental design including six different microbial inoculations and no microbial inoculation (CK), was conducted with four replicates. We inoculated the artificial soils with each bacterial species (BM: B. subtilis, SM: S. cinnamoneus). Additionally, we prepared mixed cultures, including an equal mixture of BM and SM (SB1) with the same microbial biomass as the monocultures, and three additional mixtures (SB2, SB0.5, SB0.1) with varying proportions of *B. subtilis* (**Table S1**). For the measurement of glucose mineralization in each treatment, each sample was added with a substrate solution containing glucose (0.7 mg C g^−1^ soil) and NH_4_NO_3_ (0.082 mg N g^−1^ soil), and 1×PBS buffer (0.012 mg P g^−1^ soil) to prevent any microbial nutrient limitation during the incubation period. The amount of applied glucose reflects the amount of C released from the plant roots into the soil in one week (Kallenbach et al., 2016; Su et al, 2021). The CNP ratio of the substrate was equivalent to the global mean microbial biomass ratio (60:7:1; Cleveland & Liptzin, 2007). The incubation experiment was then maintained by wetting 20% of the soil moisture with substrate solutions and microbial inoculums. The substrate solutions and microbial mixtures were mixed into the artificial soils using a sterilized spatula.

Microbial respiration (CO_2_ emission) measurement was performed using an alkali trap method. Each sample was placed inside a 225 mL glass jar containing 3 mL of 1 M NaOH and 10 mL of 0.01 M HCl which was also used for keeping the water contents of the soil. We incubated at 30°C for 14 days, and 1M NaOH was replaced new one five times (2, 4, 6, 8, and 10 days after inoculation). Exchanged 1 M NaOH was titrated with 1M HCl. Microbial respiration was calculated by subtracting non-microbial treatment (CK) from microbial treatments. At the end of the incubation experiment, the pH of each soil sample was determined by mixing 6 g of air-dried soil with 30 mL of deionized water, shaking for 30 minutes, and measuring the pH using a pH sensor (HORIBA Scientific, Japan).

### Statistical Analysis

All statistical analyses were performed using R (version 3.1.1). Tukey HSD was performed to detect differences within the treatments.

## Results and discussion

As shown in **Fig.1a**, cumulative microbial respiration was observed throughout the experiment, indicating that bacterial life strategies determine the magnitude of respiration. According to a previous study, the phyla Firmicutes and Actinobacteria could be assigned both r-strategists and K-strategists (Trivedi et al., 2013; Zhen et al., 2017; Wang et al.,2021; Bram et al., 2023). In this study, *S. cinnamoneus* (Actinobacteria) had lower mineralization rates than *B. subtilis* (Firmicutes). These differences suggest specific C dynamics driven by their contrasting life strategies. Our findings align with those reported by Guenet et al. (2011), who examined mineralization rates of six bacterial strains in artificial soils. They found that K-strategists showed delayed peaks in mineralization rates compared to r-strategists. The temporal respiration patterns further supported this distinction: *B. subtilis* respiration peaked at 2 days after inoculation, whereas *S. cinnamoneus* showed a delayed respiration peak beyond 2 days after inoculation (**Fig.1b**). Moreover, *B. subtilis* showed higher cumulative respiration per the amount of inoculated DNA than *S. cinnamoneus* (**Fig. S2**), suggesting that r-strategists have higher mineralization rates per cell than K-strategists. Our findings is consistent with a previous study showing that r-strategists require high nutritional uptake per unit cell mass to maintain their rapid growth strategy (Chen et al., 2016).

**Figure 1.**
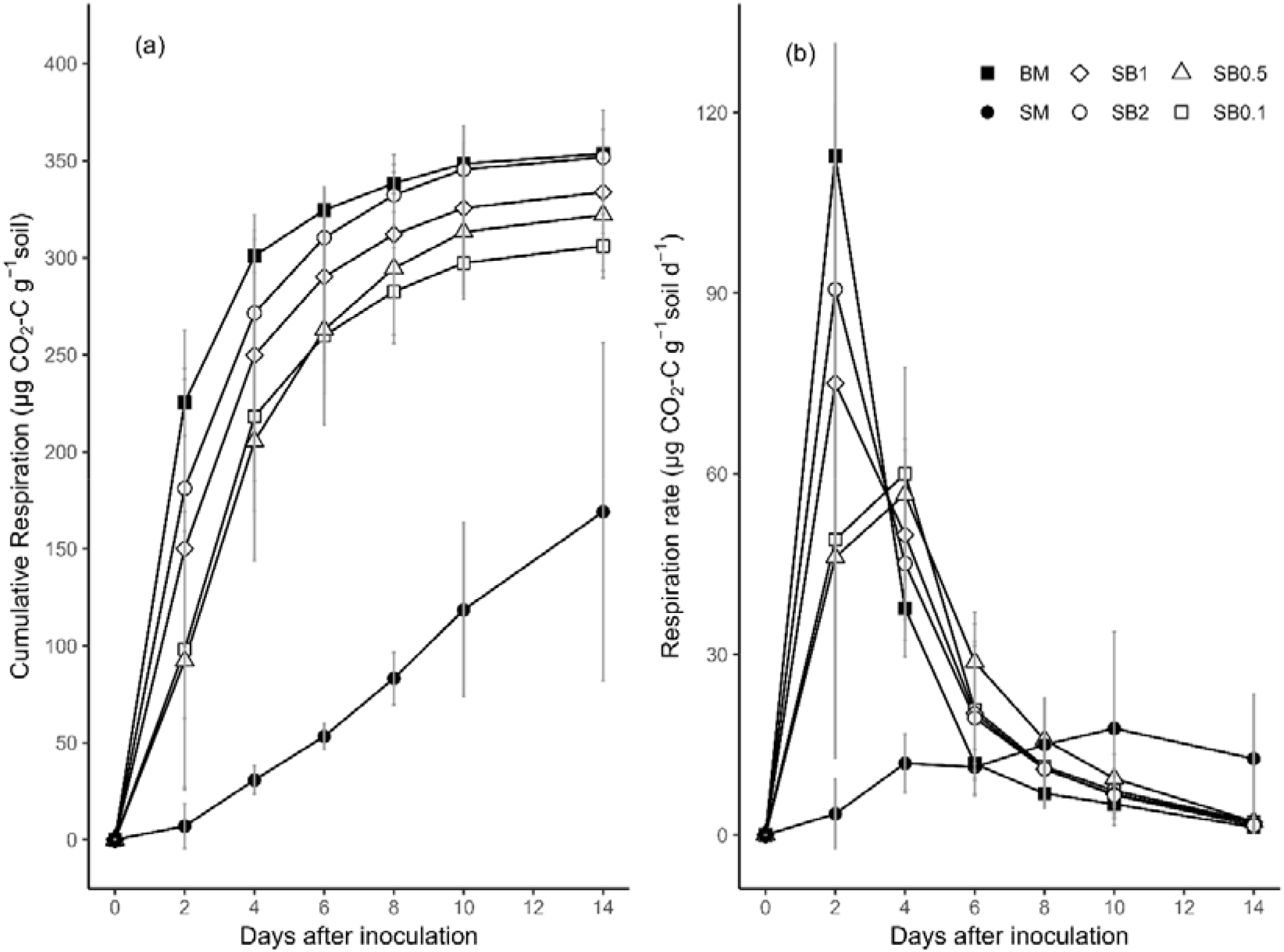
(a) Temporal cumulative respiration (μg CO□-C g−^1^ soil) and (b) microbial respiration rate (μg CO□ -C g−^1^ soil d □^1^) over 14 days of incubation. Error bars represent the standard deviation (n = 4). BM: *B. subtilis*; SM: *S*.*cinnamoneus*; SB1, SB2, SB0.5, and SB0.1: co-culture treatments with different relative abundances of *B. subtilis* (see Table S1 for details).

The cumulative respiration of co-culture treatments converged to that of *B. subtilis* monoculture treatments (**Fig.1**). This convergence may be explained by the contrasting life strategies and the possible inhibitory effects of that *B. subtilis* on the growth of *S. cinnamoneus*. Firstly, previous reports have demonstrated that r-strategists uptake glucose more rapidly and grow faster than K-strategists (Fontaine et al., 2003; Dungait et al., 2011; Wolf et al., 2013). Additionally, r-strategists have been shown to dominate over K-strategists in the presence of labile organic C such as glucose (Zhang et al., 2019). Secondly, *B. subtilis* forms biofilms under high glucose concentration and suppress competing other microbes by restricting their access to soil pore spaces (Coyte et al., 2017; Wu et al., 2019). Furthermore, *Bacillus* spp. is known as spore-forming soil bacteria and produces an antibiotic substance (Selegato et al., 2023; Lozano-Andrade et al., 2025). Previous reports have found that antibiotic substance produced by *B. subtilis*, such as surfactin and lactonase-homologous proteins, inhibit aerial hyphae development or streptomycin production in Streptomyces species (Shank et al., 2011; Hoefler et al., 2012; Schneider et al., 2012).

Our study experimentally demonstrated the relationship between the relative ratio of r/K-strategists and microbial respirations. The growth inhibition effect of *B. subtilis* on *S. cinamoneus* varied depending on the relative ratio of *B. subtilis* in the co-cultures. The respiration peak was delayed with decreasing the relative abundance of *B. subtilis* (**Fig.1b**). Furthermore, cumulative respirations showed a positive correlation with the relative abundance of *B. subtilis* in artificial soils (**Fig.S3**). These results supported previous findings that the relative abundance of r-strategists correlates positively with C mineralization rates in natural soil systems (Fierer et al., 2007; Zhang et al., 2018; Yang et al., 2021). Despite being outcompeted in co-cultures, *S. cinnamoneus* demonstrated distinctive metabolic characteristics in monoculture. K-strategists like *S. cinnamoneus* would begin to grow gradually using the intracellular stores regardless of the quantity of accessible substrates in the soil environments (Blagodatskaya et al., 2007). Notably, after the incubation period, the soils with *S*.*cinnamones* significantly decreased pH compared to other treatments (**Fig.S4**). This pH reduction is consistent with the ability of Actinobacteria to produce organic acids in the soil environments (Strzelczyk et al., 1986). Previous studies have demonstrated *Streptomyces* species produce organic acid (e.g., pyruvate, 2-oxoglutarate) into extracellular media when glucose is the primary C source, subsequently decreased medium pH (Michael et al, 1987; Madden et al, 1996).

## Conclusion

This study revealed that microbial life strategies and interspecific interactions play a critical role in shaping carbon (C) mineralization, as evidenced by experiments in an artificial soil system. *B. subtilis* exhibited rapid glucose mineralization with early respiration peaks, while *S. cinnamoneus* showed delayed respiration patterns, supporting the classification of r/K-strategist. Furthermore, co-culture experiments revealed that cumulative respiration converged to the monoculture of *B. subtilis* with cumulative respiration positively correlating with its relative abundance. These results validate artificial soil as an effective tool for elucidating microbial contributions to C dynamics, providing a foundation for understanding and predicting C cycling processes in complex natural soils. Future research should expand this approach to include more diverse microbial species and environmental conditions to better represent natural soil processes.

## Author’s Contributions

K.F., T.M., and T.H. planned and designed the research. K.F. and T.H. conducted the incubation experiments. K.F. and T.H. analyzed the data, wrote the main manuscript, and prepared the figures and tables. All authors reviewed the manuscript.

## Acknowledgments

We thank Mrs. Kondo for their kind technical support in the incubation experiments. This work was financially supported by the JSPS KAKENHI Grant Number (JP23K14056), and the JGC Saneyoshi Scholarship Foundation (2302)

## Competing Interests Statement

The authors declare no competing interests.

## Notes

### Competing Interest Statement

The authors have declared no competing interest.

## Reference

1. L. Aliperti, A. A. Aptekmann, Gonzalo Farfañuk, L. L. Couso, A. Soler-Bistué, and I. E. S. Anchez, “r/K selection of GC content in prokaryotes,” 2023, doi: 10.1111/1462-2920.16511.

2. Z. Bai, H. Xie, J. Kao-Kniffin, B. Chen, P. Shao, and C. Liang, “Shifts in microbial trophic strategy explain different temperature sensitivity of CO2 flux under constant and diurnally varying temperature regimes,” FEMS Microbiology Ecology, vol. 93, no. 5, May 2017, doi: 10.1093/FEMSEC/FIX063.

3. C. Baker, J. De, K. Schneider, and C. Keith Schneider, “Escherichia coli O157 survival in liquid culture and artificial soil microcosms with variable pH, humic acid and clay content,” 2020, doi: 10.1111/jam.14775.

4. E. Blagodatskaya, S. Blagodatsky, M. Dorodnikov, and Y. Kuzyakov, “Elevated atmospheric CO2 increases microbial growth rates in soil: results of three CO2 enrichment experiments,” Global Change Biology, vol. 16, no. 2, pp. 836–848, Feb. 2010, doi: 10.1111/J.1365-2486.2009.02006.X.

5. E. v. Blagodatskaya, S. A. Blagodatsky, T. H. Anderson, and Y. Kuzyakov, “Priming effects in Chernozem induced by glucose and N in relation to microbial growth strategies,” Applied Soil Ecology, vol. 37, no. 1–2, pp. 95–105, Oct. 2007, doi: 10.1016/J.APSOIL.2007.05.002.

6. H. Chen et al., “Microbial respiratory thermal adaptation is regulated by r-/K-strategy dominance,” Ecology Letters, vol. 25, no. 11. John Wiley and Sons Inc, pp. 2489–2499, Nov. 01, 2022. doi: 10.1111/ele.14106.

7. Y. Chen et al., “Large amounts of easily decomposable carbon stored in subtropical forest subsoil are associated with r-strategy-dominated soil microbes,” Soil Biology and Biochemistry, vol. 95, pp. 233–242, Apr. 2016, doi: 10.1016/J.SOILBIO.2016.01.004.

8. Y. Cheng and W. Wan, “Elevated salinity decreases soil ecosystem multifunctionality by shifting the bacterial community from K-to r-selected living strategy,” Land Degradation and Development, Feb. 2022, doi: 10.1002/ldr.4519

9. C. C. Cleveland and D. Liptzin, “C:N:P stoichiometry in soil: Is there a ‘Redfield ratio’ for the microbial biomass?,” Biogeochemistry, vol. 85, no. 3, pp. 235–252, Sep. 2007, doi: 10.1007/s10533-007-9132-0.

10. K. Z. Coyte, H. Tabuteau, E. A. Gaffney, K. R. Fostera, and W. M. Durham, “Microbial competition in porous environments can select against rapid biofilm growth,” Proceedings of the National Academy of Sciences of the United States of America, vol. 114, no. 2, pp. E161–E170, Jan. 2017, doi: 10.1073/pnas.1525228113.

11. del Valle, X. Gao, T. A. Ghezzehei, J. J. Silberg, and C. A. Masiello, “Artificial Soils Reveal Individual Factor Controls on Microbial Processes,” mSystems, vol. 7, no. 4, Aug. 2022, doi: 10.1128/msystems.00301-22

12. T. O. Delmont et al., “Microbial community development and unseen diversity recovery in inoculated sterile soil,” Biology and Fertility of Soils, vol. 50, no. 7, pp. 1069–1076, Sep. 2014, doi: 10.1007/S00374-014-0925-8/FIGURES/8.

13. L. A. Domeignoz-Horta, G. Pold, X. J. A. Liu, S. D. Frey, J. M. Melillo, and K. M. DeAngelis, “Microbial diversity drives carbon use efficiency in a model soil,” Nature Communications, vol. 11, no. 1, Dec. 2020, doi: 10.1038/s41467-020-17502-z

14. J. A. J. Dungait et al., “Variable responses of the soil microbial biomass to trace concentrations of 13C-labelled glucose, using 13C-PLFA analysis,” European Journal of Soil Science, vol. 62, no. 1, pp. 117–126, Feb. 2011, doi: 10.1111/j.1365-2389.2010.01321.x.

15. R. J. Ellis, “Artificial soil microcosms: A tool for studying microbial autecology under controlled conditions,” Journal of Microbiological Methods, vol. 56, no. 2, pp. 287–290, 2004, doi: 10.1016/j.mimet.2003.10.005.

16. N. Fierer, M. A. Bradford, and R. B. Jackson, “Toward an ecological classification of soil bacteria,” Ecology, vol. 88, no. 6, pp. 1354–1364, Jun. 2007, doi: 10.1890/05-1839.

17. S. Fontaine, A. Mariotti, and L. Abbadie, “The priming effect of organic matter: a question of microbial competition?,” Soil Biology and Biochemistry, vol. 35, no. 6, pp. 837–843, Jun. 2003, doi: 10.1016/S0038-0717(03)00123-8.

18. B. Guenet, J. Leloup, C. Hartmann, S. Barot, and L. Abbadie, “A new protocol for an artificial soil to analyse soil microbiological processes,” Applied Soil Ecology, vol. 48, no. 2, pp. 243–246, Jun. 2011, doi: 10.1016/j.apsoil.2011.04.002

19. B. C. Hoefler, K. v. Gorzelnik, J. Y. Yang, N. Hendricks, P. C. Dorrestein, and P. D. Straight, “Enzymatic resistance to the lipopeptide surfactin as identified through imaging mass spectrometry of bacterial competition,” Proceedings of the National Academy of Sciences of the United States of America, vol. 109, no. 32, pp. 13082–13087, Aug. 2012, doi: 10.1073/pnas.1205586109.

20. P. Hu et al., “Linking bacterial life strategies with soil organic matter accrual by karst vegetation restoration,” Soil Biology and Biochemistry, vol. 177, p. 108925, Feb. 2023, doi: 10.1016/J.SOILBIO.2022.108925.

21. H. Iven, T. W. N. Walker, and M. Anthony, “Biotic Interactions in Soil are Underestimated Drivers of Microbial Carbon Use Efficiency,” vol. 80, p. 13, 2023, doi: 10.1007/s00284-022-02979-2.

22. M. Jiang et al., “Home□based microbial solution to boost crop growth in low□fertility soil,” New Phytologist, May 2023, doi: 10.1111/nph.18943.

23. C. M. Kallenbach, S. D. Frey, and A. S. Grandy, “Direct evidence for microbial-derived soil organic matter formation and its ecophysiological controls,” Nature Communications, vol. 7, Nov. 2016, doi: 10.1038/ncomms13630.

24. J. T. Lennon, Z. T. Aanderud, B. K. Lehmkuhl, and D. R. Schoolmaster, “Mapping the niche space of soil microorganisms using taxonomy and traits,” Ecology, vol. 93, no. 8, pp. 1867–1879, Aug. 2012, doi: 10.1890/11-1745.1.

25. K. Li, M. J. DiLegge, I. S. Minas, A. Hamm, D. Manter, and J. M. Vivanco, “Soil sterilization leads to re-colonization of a healthier rhizosphere microbiome,” Rhizosphere, vol. 12, Dec. 2019, doi: 10.1016/j.rhisph.2019.100176.

26. J. Li, L. Zhao, C. Song, C. He, H. Bian, and L. Sheng, “Forest swamp succession alters organic carbon composition and survival strategies of soil microbial communities,” Science of the Total Environment, vol. 904, Dec. 2023, doi: 10.1016/j.scitotenv.2023.166742.

27. C. Liang, W. Amelung, J. Lehmann, and M. Kästner, “Quantitative assessment of microbial necromass contribution to soil organic matter,” Global Change Biology, vol. 25, no. 11, pp. 3578–3590, Nov. 2019, doi: 10.1111/GCB.14781.

28. C. N. Lozano-Andrade et al., “Surfactin facilitates establishment of Bacillus subtilis in synthetic communities,” The ISME Journal, vol. 19, no. 1, Jan. 2025, doi: 10.1093/ISMEJO/WRAF013.

29. T. Madden, J. M. Ward, and A. P. Ison, “Organic acid excretion by Streptomyces lividans TK24 during growth on defined carbon and nitrogen sources,” 1996.

30. Y. Maeda et al., “Invention of artificial rice field soil: A tool to study the effect of soil components on the activity and community of microorganisms involved in anaerobic organic matter decomposition,” Microbes and Environments, vol. 35, no. 4, pp. 1–10, 2020, doi: 10.1264/jsme2.ME20093.

31. G. J. Michaelson, C. L. Ping, and G. A. Mitchell, “Correlation of Mehlich 3, Bray 1, and ammonium acetate extractable P, K, Ca, and Mg for alaska agricultural soils,” Communications in Soil Science and Plant Analysis, vol. 18, no. 9, pp. 1003–1015, Sep. 1987, doi: 10.1080/00103628709367877.

32. L. Nazaries, J. C. Murrell, P. Millard, L. Baggs, and B. K. Singh, “Methane, microbes and models: Fundamental understanding of the soil methane cycle for future predictions,” Environmental Microbiology, vol. 15, no. 9, pp. 2395–2417, Sep. 2013, doi: 10.1111/1462-2920.12149/SUPPINFO.

33. G. W. Nicol et al., “Mineral vs. Organic Amendments: Microbial Community Structure, Activity and Abundance of Agriculturally Relevant Microbes Are Driven by Long-Term Fertilization Strategies,” 2016, doi: 10.3389/fmicb.2016.01446

34. N. Nonthakaew, W. Panbangred, W. Songnuan, and B. Intra, “Plant growth-promoting properties of Streptomyces spp. isolates and their impact on mung bean plantlets’ rhizosphere microbiome,” Frontiers in Microbiology, vol. 13, p. 967415, Aug. 2022, doi: 10.3389/FMICB.2022.967415/BIBTEX.

35. E. R. Pianka, “On r- and K-Selection.”, American Naturalist 104:592–597.

36. F. Rakhsh, A. Golchin, A. B. al Agha, and P. N. Nelson, “Mineralization of organic carbon and formation of microbial biomass in soil: Effects of clay content and composition and the mechanisms involved,” 2020, doi: 10.1016/j.soilbio.2020.108036.

37. K. Sawada, S. Funakawa, and T. Kosaki, “Soil microorganisms have a threshold concentration of glucose to increase the ratio of respiration to assimilation,” Soil Science & Plant Nutrition, vol. 54, no. 2, pp. 216–223, Apr. 2008, doi: 10.1111/J.1747-0765.2007.00235.X.

38. K. Saxena, M. Kumar, H. Chakdar, N. Anuroopa, and D. J. Bagyaraj, “Bacillus species in soil as a natural resource for plant health and nutrition,” Journal of Applied Microbiology, vol. 128, no. 6, pp. 1583–1594, Jun. 2020, doi: 10.1111/JAM.14506.

39. J. Schneider, A. Yepes, J. C. Garcia-Betancur, I. Westedt, B. Mielich, and D. López, “Streptomycin-induced expression in Bacillus subtilis of YtnP, a lactonase-homologous protein that inhibits development and streptomycin production in Streptomyces griseus,” Applied and Environmental Microbiology, vol. 78, no. 2, pp. 599–603, Jan. 2012, doi: 10.1128/AEM.06992-11.

40. D. M. Selegato and I. Castro-Gamboa, “Enhancing chemical and biological diversity by co-cultivation,” Frontiers in Microbiology, vol. 14. Frontiers Media S.A., Feb. 01, 2023. doi: 10.3389/fmicb.2023.1117559

41. Q. Shang and Y. Liu, “Differences in Soil CO2 Efflux and Microbial Community Composition Among Slope Aspects in a Mountain Oak Forest,” Forests, vol. 15, no. 10, Oct. 2024, doi: 10.3390/f15101810.

42. E. A. Shank, V. Klepac-Ceraj, L. Collado-Torres, G. E. Powers, R. Losick, and R. Kolter, “Interspecies interactions that result in Bacillus subtilis forming biofilms are mediated mainly by members of its own genus,” Proceedings of the National Academy of Sciences of the United States of America, vol. 108, no. 48, Nov. 2011, doi: 10.1073/PNAS.1103630108.

43. B. W. G. Stone et al., “Life history strategies among soil bacteria—dichotomy for few, continuum for many,” The ISME Journal 2023 17:4, vol. 17, no. 4, pp. 611–619, Feb. 2023, doi: 10.1038/s41396-022-01354-0.

44. P. D. Straight, J. M. Willey, and R. Kolter, “Interactions between Streptomyces coelicolor and Bacillus subtilis: Role of surfactants in raising aerial structures,” Journal of Bacteriology, vol. 188, no. 13, pp. 4918–4925, Jul. 2006, doi: 10.1128/JB.00162-06/ASSET/E3CB3E6C-30FE-487A-BF88-FC31CE50E5A6/ASSETS/GRAPHIC/ZJB0130658410003.JPEG.

45. E. Strzelczyk, “Organic acids production by Streptomyces spp. isolated from soil, rhizosphere and mycorrhizosphere of pine (Pinus sylvestris L.).”

46. M. Su et al., “Phosphorus deficiency in soils with red color: Insights from the interactions between minerals and microorganisms,” Geoderma, vol. 404, Dec. 2021, doi: 10.1016/j.geoderma.2021.115311.

47. Y. Sun, C. Wang, J. Yang, J. Liao, H. Y. H. Chen, and H. Ruan, “Elevated CO2 shifts soil microbial communities from K-to r-strategists,” Global Ecology and Biogeography, vol. 30, no. 5, pp. 961–972, May 2021, doi: 10.1111/geb.13281.

48. V. Torsvik and L. Øvreås, “Microbial diversity and function in soil: from genes to ecosystems,” Current Opinion in Microbiology, vol. 5, no. 3, pp. 240–245, Jun. 2002, doi: 10.1016/S1369-5274(02)00324-7.

49. P. Trivedi, I. C. Anderson, and B. K. Singh, “Microbial modulators of soil carbon storage: Integrating genomic and metabolic knowledge for global prediction,” Trends in Microbiology, vol. 21, no. 12, pp. 641–651, Dec. 2013, doi: 10.1016/J.TIM.2013.09.005.

50. H. Wang et al., “Eight years of manure fertilization favor copiotrophic traits in paddy soil microbiomes,” European Journal of Soil Biology, vol. 106, p. 103352, Sep. 2021, doi: 10.1016/J.EJSOBI.2021.103352.

51. X. Wang et al., “Mineral composition controls the stabilization of microbially derived carbon and nitrogen in soils: Insights from an isotope tracing model,” Global Change Biology, vol. 30, no. 1, Jan. 2024, doi: 10.1111/gcb.17156.

52. X. Wang et al., “Organic amendments drive shifts in microbial community structure and keystone taxa which increase C mineralization across aggregate size classes,” Soil Biology and Biochemistry, vol. 153, p. 108062, Feb. 2021, doi: 10.1016/J.SOILBIO.2020.108062

53. S. Wertz et al., “Early-stage bacterial colonization between a sterilized remoulded soil clod and natural soil aggregates of the same soil,” Soil Biology and Biochemistry, vol. 39, no. 12, pp. 3127–3137, Dec. 2007, doi: 10.1016/J.SOILBIO.2007.07.005.

54. B. Wolf, M. Vos, W. de Boer, and G. A. Kowalchuk, “Impact of matric potential and pore size distribution on growth dynamics of filamentous and non-filamentous soil bacteria,” PLoS ONE, vol. 8, no. 12, Dec. 2013, doi: 10.1371/journal.pone.0083661.

55. D. C. Wolf, T. H. Dao, H. D. Scott, and T. L. Lavy, “Influence of Sterilization Methods on Selected Soil Microbiological, Physical, and Chemical Properties.”

56. Y. Wu et al., “Soil biofilm formation enhances microbial community diversity and metabolic activity,” Environment International, vol. 132, p. 105116, Nov. 2019, doi: 10.1016/J.ENVINT.2019.105116

57. B. Yan, N. Liu, M. Liu, X. Du, F. Shang, and Y. Huang, “Soil actinobacteria tend to have neutral interactions with other co-occurring microorganisms, especially under oligotrophic conditions,” Environmental Microbiology, vol. 23, no. 8, pp. 4126–4140, Aug. 2021, doi: 10.1111/1462-2920.15483.

58. Y. Yang et al., “Negative effects of multiple global change factors on soil microbial diversity,” Soil Biology and Biochemistry, vol. 156, p. 108229, May 2021, doi: 10.1016/J.SOILBIO.2021.108229.

59. Z. Zhang, W. Wang, J. Qi, H. Zhang, F. Tao, and R. Zhang, “Priming effects of soil organic matter decomposition with addition of different carbon substrates,” Journal of Soils and Sediments, vol. 19, no. 3, pp. 1171–1178, Mar. 2019, doi: 10.1007/s11368-018-2103-3.

60. X. Zhang et al., “Tree species mixture inhibits soil organic carbon mineralization accompanied by decreased r-selected bacteria,” Plant and Soil, vol. 431, no. 1–2, pp. 203–216, Oct. 2018, doi: 10.1007/S11104-018-3755-X/FIGURES/5.

61. W. Zheng et al., “The response patterns of r- and K-strategist bacteria to long-term organic and inorganic fertilization regimes within the microbial food web are closely linked to rice production,” Science of the Total Environment, vol. 942, Sep. 2024, doi: 10.1016/j.scitotenv.2024.173681.

