## Supplemental Figures for "Artificial soil systems: A tool for investigating microbial life strategy effects on substrate mineralization"

Number of Figures: 4

Number of Tables: 1

Keywords: artificial soil, C mineralization, r/K strategist, interspecific interaction

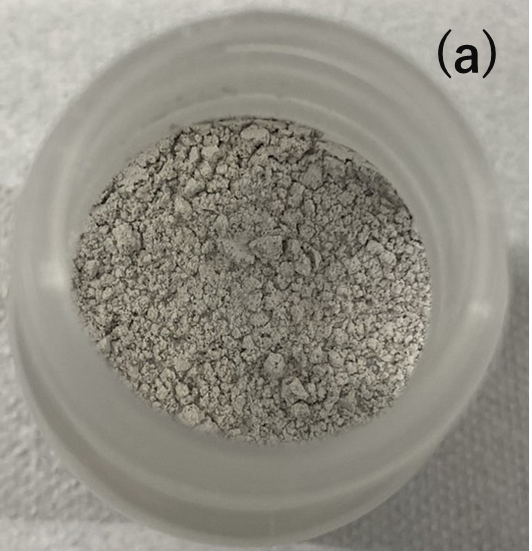
　 　
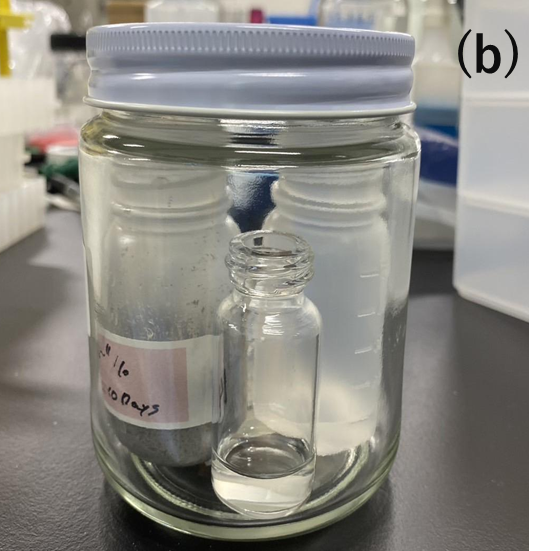

Figure S1. Photographs of (a) artificial soil mixture and (b) an alkali trap setup. The alkali trap consists of three bottles: a bottle containing the artificial soils, a glass vial with 1 M NaOH, and another bottle with 0.01 M HCl to maintain the soil moisture.

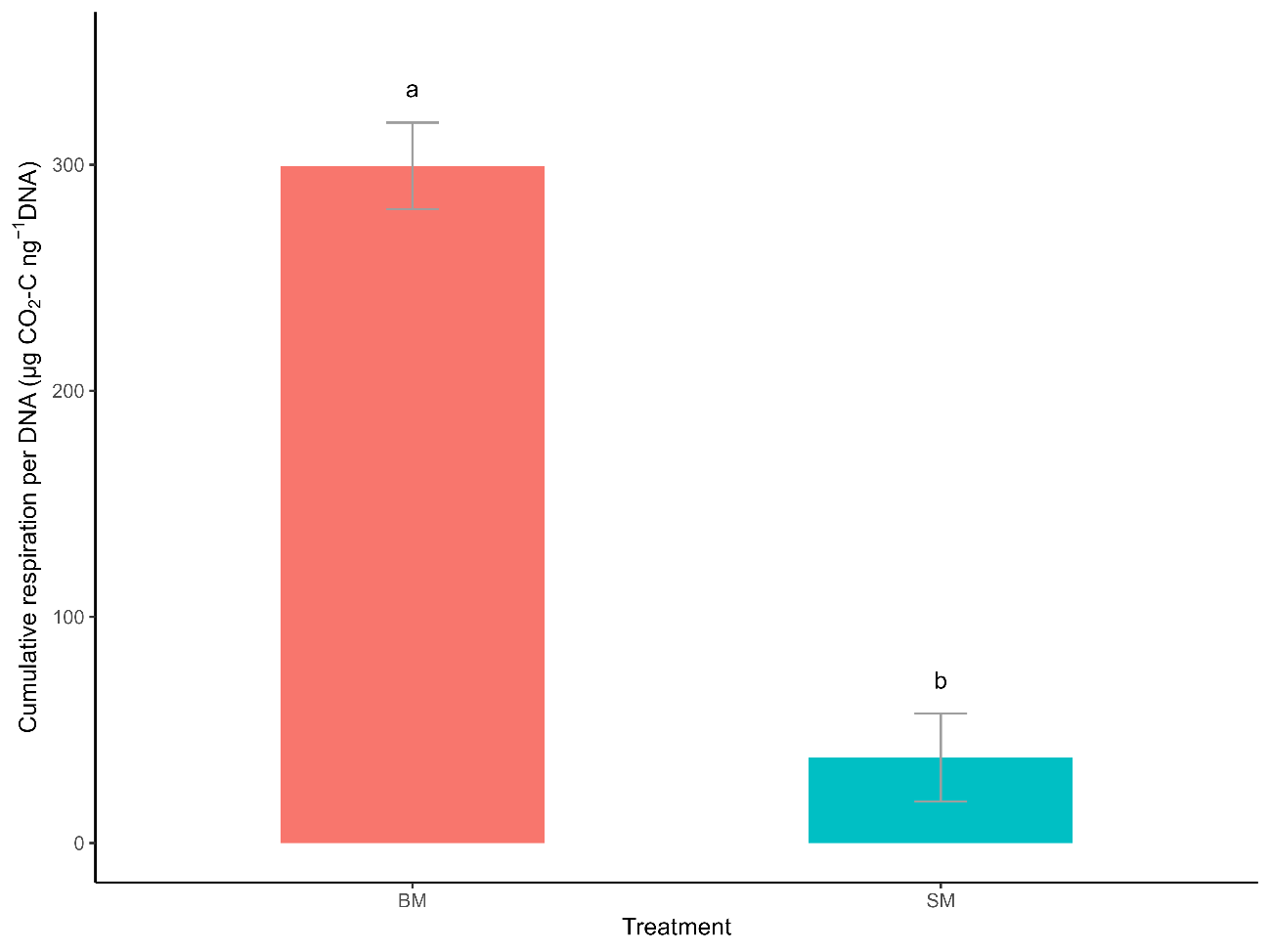

Figure S2. Cumulative respiration per the amount of inoculated DNA in monoculture treatments (BM: B. subtilis, SM: S. cinnamoneus). Error bars represent the standard deviation (n = 4). Different letters indicate significant differences (p < 0.05).

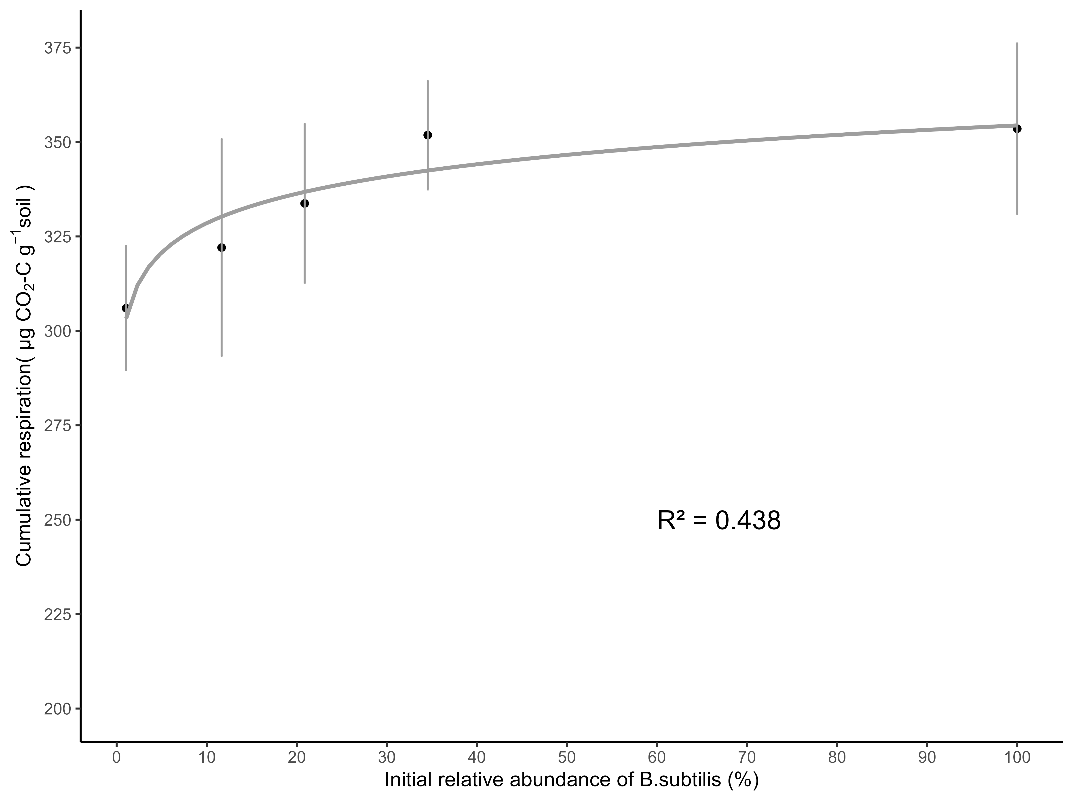

Figure S3. Relationship between cumulative respiration (μg CO₂-C g^−1^ soil) and the relative abundance of inoculated *B. subtilis* (%). Error bars represent the standard deviation (n = 4). The relationship was fitted using a logarithmic model (R² = 0.438).

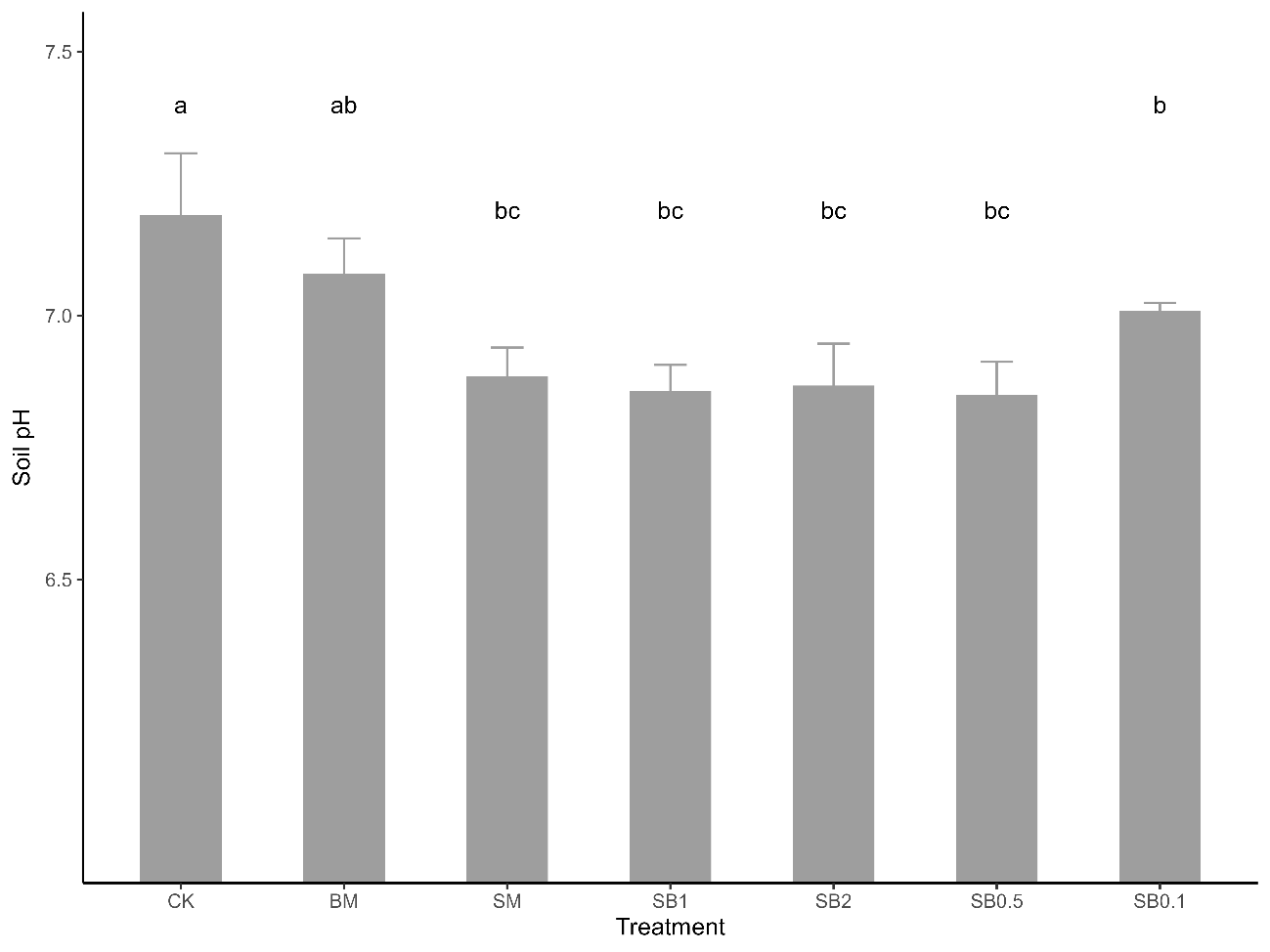

Figure S4. Soil pH at the end of incubation. Error bars represent the standard deviation (n = 4). Different letters indicate significant differences (p < 0.05). CK: control; BM: B. subtilis; SM: S.cinnamoneus; SB1, SB2, SB0.5, and SB0.1: co-culture treatments with different relative abundances of B. subtilis (see Table S1 for details).
